# CoDNaS-Q: a database of conformational diversity of the native state of proteins with quaternary structure

**DOI:** 10.1101/2022.02.15.480082

**Authors:** Nahuel Escobedo, Ronaldo Romario Tunque Cahui, Gastón Caruso, Emilio García Ríos, Layla Hirsh, Alexander Miguel Monzon, Gustavo Parisi, Nicolas Palopoli

## Abstract

**Summary:** A collection of conformers that exist in a dynamical equilibrium defines the native state of a protein. The structural differences between them describe their conformational diversity, a defining characteristic of the protein with an essential role in multiple cellular processes. Since most proteins carry out their functions by assembling into complexes, we have developed CoDNaS-Q, the first online resource to explore conformational diversity in homooligomeric proteins. It features a curated collection of redundant protein structures with known quaternary structure. CoDNaS-Q integrates relevant annotations that allow researchers to identify and explore the extent and possible reasons of conformational diversity in homooligomeric protein complexes.

**Availability and implementation:** CoDNaS-Q is freely accessible at http://ufq.unq.edu.ar/codnasq/. The data can be retrieved from the website. The source code of the database can be downloaded from https://github.com/SfrRonaldo/codnas-q.

## Introduction

The native state of a protein is represented by an ensemble of alternative conformers in equilibrium. Structural differences between these conformers define the extent of the protein’s conformational diversity, from local loop rearrangements to large displacements of entire subunits. These conformational changes are usually of biological relevance, accounting for diverse functional mechanisms (Monzon et al., 2017) and related with processes such as enzyme catalysis (Henzler-Wildman et al., 2007), promiscuity (James and Tawfik, 2003), binding specificity (Yogurtcu et al., 2008), allosterism (Changeux, 2012) and cell signaling (Nussinov and Ma, 2012).

Most proteins carry out their functions by assembling into complexes, forming obligate or non-obligate (permanent or transient) interactions between two or more polypeptide chains (Nooren and Thornton, 2003). This is reflected in structural databases, where more than half of the known structures are multimeric, generally forming homooligomers (Bertoni et al., 2017); (Levy et al., 2006). Conformational diversity is functionally relevant at the quaternary structure (QS) level too. It has been shown that the conformational diversity of proteins upon complex formation is strongly dependent on the flexibility of their free monomers (Marsh et al., 2012). The degree of protein flexibility is also associated with different quaternary topologies, reflecting the structural constraints on evolutionary history imposed by characteristic solvent accessible surface areas and interface contacts between subunits (Marsh and Teichmann, 2014). Addressing conformational diversity at all structural levels appears essential in pursuing a better understanding of protein function.

Our group has previously developed CoDNaS, a database of Conformational Diversity on the Native State of proteins (Monzon et al., 2016). It has allowed us to explore the structural and functional implications of conformational diversity of individual protein chains. (We recently published CoDNaS-RNA with the similar goal of addressing conformational diversity of single RNA chains (Buitrón et al., 2021)). In this work we extend our approach by presenting CoDNaS-Q, a database of conformational diversity of the native state of proteins with QS. Each entry in CoDNaS-Q presents a cluster of known structures of the same complex available from the Protein Data Bank (PDB) (Berman et al., 2000), along with all-vs-all structural comparisons and annotations to facilitate the inspection of its conformational diversity. CoDNaS-Q is currently limited to homooligomers, following their overrepresentation in the PDB.

## Dataset

Experimental-based evidence of conformational diversity comes from the analysis of known protein structures in the Protein Data Bank (wwPDB consortium, 2019). A collection of crystallographically determined structures of a protein can be interpreted as snapshots of its native conformational ensemble. Thereby, conformational diversity can be studied using redundant structures of the same protein obtained in different experimental conditions (with or without ligands, varying pH or temperature, etc). We used sequence clusters of all individual chains from the PDB obtained with BLASTClust at 95% sequence identity, and only kept clusters with two or more chains from two or more PDB files (see Figure 1, panel A for an overview of the development scheme). Each cluster can be interpreted as a set of native conformers of a homooligomeric protein.

**Figure 1.**
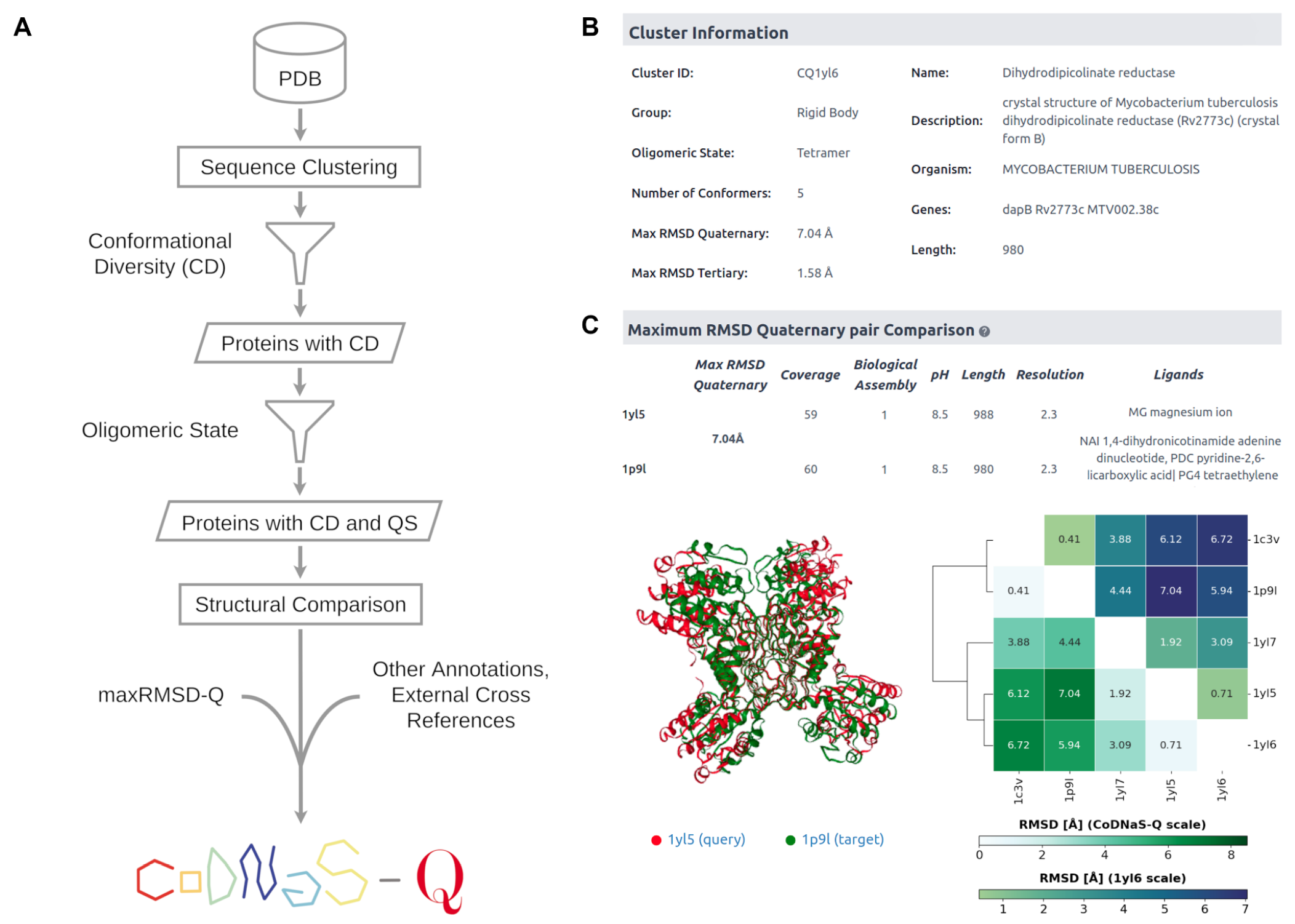
**Panel A:** CoDNaS-Q development scheme. **Panel B:** Cluster Information section from a sample entry in CoDNaS-Q. The table presents general data about the cluster and selected annotations from a representative conformer. **Panel C:** Comparison of conformers from a sample entry in CoDNaS-Q. The table highlights data from the pair of conformers with maximum RMSD at the quaternary level that is useful to understand the extent and possible reasons of the observed conformational diversity in the cluster. An interactive window provides the structural superposition of the maximum RMSD conformers. A dendrogram and a heatmap of the RMSD-based hierarchical clustering of the available conformers facilitates inspection of structural differences between any pair of conformers.

The preliminary set of clusters was filtered to retain only those with a reliably assigned oligomeric state. To this end, the conformers with a biologically relevant QS were selected with QSbio, a method to annotate the QS of biological assemblies with high confidence by combining prediction tools and databases, and thus achieve accuracy close to manual curation (Dey et al., 2018). CoDNaS-Q is limited to conformers with two or more subunits and High or Very High confidence on QSbio annotations of their QS. We also discard entries with ambiguous QS assignments. The first release of CoDNaS-Q comprises 3649 clusters and 18790 conformers. In the near future we plan to update CoDNaS-Q with more clusters comprising homo- and hetero-oligomers, with an API REST service for data exchange with users and other data resources.

The RMSD (Root Mean Square Deviation) is a common metric used in structural biology to measure the dissimilarity between two biological structures that also provides a valid estimation of a protein’s conformational diversity. CoDNaS-Q uses the RMSD calculated with TopMatch (Sippl and Wiederstein, 2012) to compare multimeric complexes and individual chains. We compute the Cα-RMSD between all pairs of multimeric conformers within every cluster and identify the maximum RMSD at the quaternary level (maxRMSD-Q) to quantify the degree of conformational diversity in the ensemble. TopMatch also provides other parameters useful to characterize an ensemble of conformers such as typical distance error, alignment coverage, number of structurally equivalent residue pairs, etc. Each cluster is annotated with additional data like the maximum RMSD between any two individual chains (maxRMSD-T) and features extracted from the PDB (source organism, experimental method, pH and temperature, ligands, etc), UniProt (cross-referenced gene name and entry ID), and Pfam (family ID).

## Website

The website allows users to search CoDNaS-Q by Cluster ID, PDB code, name and description of an entry, source organism, related UniProt accession ID, type of movement and oligomeric state, among other criteria.

At the top of each CoDNaS-Q entry, the Cluster Information section (Figure 1, panel B) presents general information about the structural properties and the conformational diversity related with the cluster, along with selected annotations from a representative conformer. Below, a table compares relevant features from the maximum pair of conformers, i.e. those with the maximum RMSD value between them (Figure 1, panel C). It also includes the interactive superposition of the structures (available for download). A summary of the hierarchical clustering of conformers based on their pairwise RMSD values is shown as a dendrogram, with coloring schemes that reflect the extent of conformational diversity among conformers and in comparison with the entire database. The bottom of the page displays a sortable list of all conformers in the cluster and their structural properties. Clicking on a row provides extended information about the conformer, cross-referenced from other databases, together with its 3D structure. Two or more conformers can be selected to access a dedicated page to compare all possible pairs among the selected conformers.

## Conclusions

CoDNaS-Q is a unique resource that helps the community to learn more about the structural nature of protein complexes and their conformational diversity patterns. It helps us to understand the complexity of studying the QS in proteins of biological interest, and raises questions about the dynamics that a protein can present in its native state, as well as the factors that condition this ensemble. It provides valuable insights about the extension of quaternary conformational diversity and its relationship with protein function. We think CoDNaS-Q can be an essential resource for both computational biologists and life science researchers alike.

## Acknowledgements

The authors would like to thank Martin Gonzalez Buitrón for his help in designing and reviewing CoDNaS-Q.

